# Culture-Free Rapid Phenotypic Antimicrobial Susceptibility Testing for Helicobacter pylori Based on Fluorescence Rapid On-Site Evaluation Technology: A Preliminary Study

**DOI:** 10.64898/2026.07.06.736681

**Authors:** Binghui Li, Lin Zhang, Yanhong Hou, Kai Wu, Jiafeng Han, Junyu Liu, Jing Zhang, Mi Yang

## Abstract

**Background:** Phenotypic antibiotic susceptibility testing (AST) for Helicobacter pylori (H. pylori) has relied on bacterial culture for three decades, requiring 5–7 days to yield results. Genotypic rapid tests can only detect known resistance mutations and fail to reliably identify amoxicillin resistance. To our knowledge, no culture-free rapid phenotypic AST method for H. pylori has been previously reported.

**Methods:** We developed a phenotypic AST method based on fluorescence rapid on-site evaluation (ROSE) technology that completely bypasses bacterial culture. Gastric mucosal biopsy specimens from 40 H. pylori-positive patients were homogenized and co-incubated with an acridine orange/ethidium bromide (AO/EB)-based viability staining reagent and three first-line antibiotics—amoxicillin, clarithromycin, and levofloxacin—at concentrations corresponding to the European Committee on Antimicrobial Susceptibility Testing (EUCAST) breakpoints for H. pylori, at 37 °C for 1 hour. Fluorescence intensity was measured using a microplate reader. A reduction in fluorescence relative to an antibiotic-free control indicated susceptibility, whereas no significant reduction indicated resistance. Conventional culture-based AST (E-test) served as the reference method. The overall concordance rate, sensitivity, specificity, and Cohen’s kappa coefficient were calculated.

**Results:** Fourteen of the 40 samples had unsuccessful culture and were excluded, leaving 26 samples for statistical analysis of each antibiotic. The overall concordance rates between the ROSE method and culture-based AST were 84.6% (22/26) for amoxicillin, 76.9% (20/26) for levofloxacin, and 69.2% (18/26) for clarithromycin. Cohen’s kappa coefficients indicated moderate agreement for all three antibiotics (κ = 0.523, 0.539, and 0.412, respectively). Unlike genotypic methods, the ROSE method successfully assessed amoxicillin susceptibility in all 40 patients—a critical first-line antibiotic for which no reliable genetic resistance marker currently exists. The turnaround time was approximately 1 hour (55–65 minutes), compared with 5–7 days for culture-based methods; preliminary estimates indicated a cost reduction of approximately 3,000–5,000 Chinese yuan (CNY) per patient, mainly attributable to the elimination of culture media, prolonged incubation, and repeat clinic visits.

**Conclusions:** This study reports, for the first time, a culture-free 1-hour phenotypic AST for H. pylori. The method enables same-day, susceptibility-guided treatment decisions, addressing an unmet clinical need spanning three decades. Algorithm optimization and a prospective randomized controlled trial are currently underway to further improve diagnostic accuracy and validate clinical utility.

## 1. Introduction

Helicobacter pylori (H. pylori) infection affects approximately half of the world’s population and is a major etiological factor for gastric cancer^**Error! Unknown switch argument**.^. Eradication therapy is increasingly challenged by antibiotic resistance: clarithromycin resistance already exceeds 15% in many regions, and metronidazole resistance continues to rise worldwide^**Error! Unknown switch argument**.^. Current guidelines recommend susceptibility-guided individualized therapy once the resistance rate exceeds 15%; however, conventional phenotypic susceptibility testing requires a 5–7-day culture cycle, which delays treatment decisions and forces clinicians to rely on empirical regimens^**Error! Unknown switch argument**.^.

Genotypic rapid tests (PCR, next-generation sequencing) have emerged as alternatives but carry fundamental limitations: they detect only known resistance mutations; they cannot reliably identify amoxicillin resistance—a key first-line antibiotic with no established genetic resistance marker; and they miss emerging or complex resistance mechanisms^**Error! Unknown switch argument**.^. What the clinic truly requires is a rapid phenotypic assay that combines the clinical certainty of phenotypic testing with the speed of genotypic methods.

To our knowledge, no culture-free rapid phenotypic susceptibility method that operates directly on clinical biopsy specimens has been reported for H. pylori. Here, we report for the first time a culture-free rapid phenotypic susceptibility assay based on fluorescence rapid on-site evaluation technology that can be performed directly on gastric mucosal biopsy specimens without culture. The conception, development, and preliminary validation of this method were carried out independently by our research team.

## 2. Methods

### 2.1 Study population

Between January 2026 and June 2026, we prospectively enrolled 40 consecutive H. pylori-positive adult patients undergoing esophagogastroduodenoscopy at the Eighth Medical Center of the Chinese PLA General Hospital. The inclusion criteria were: (1) H. pylori positivity confirmed by urea breath test (UBT) or rapid urease test (RUT) within 2 weeks before enrollment; (2) no history of H. pylori eradication therapy or any systemic antibiotic exposure (including for non-eradication purposes) within 4 weeks; and (3) no use of proton pump inhibitors within 2 weeks. All participants provided written informed consent. The study protocol was approved by the Institutional Review Board of the Eighth Medical Center of the Chinese PLA General Hospital (approval no. [S-2026-003-01]) and registered at [ClinicalTrials.gov/ChiCTR: ChiCTR2600125137].

### 2.2 ROSE-based rapid phenotypic susceptibility testing

#### 2.2.1 Specimen processing

Two antral biopsy specimens were obtained from each patient: one for the fluorescence ROSE assay and one for conventional culture-based susceptibility testing. The specimen for the ROSE assay was immediately placed in 1 mL of sterile phosphate-buffered saline (PBS) and ground with a sterile pestle for 30 s to release bacteria from the mucosal tissue.

#### 2.2.2 Antibiotic exposure

The prepared bacterial homogenate was divided into four 200-μL aliquots: one served as an antibiotic-free blank control, and each of the remaining three was supplemented with one antibiotic. The final exposure concentrations were set according to the EUCAST susceptibility breakpoint MICs for H. pylori—amoxicillin 0.125 μg/mL, clarithromycin 0.125 μg/mL, and levofloxacin 1.0 μg/mL—and all samples were incubated at 37 °C under microaerobic conditions (5% O□, 10% CO□, 85% N□) for 1 h^**Error! Unknown switch argument**.^. The EUCAST breakpoint concentrations were used to set the antibiotic exposure concentrations for the ROSE assay, whereas E-test interpretation followed the standard MIC breakpoints of EUCAST v16.1; the two constitute reference systems used for different purposes.

#### 2.2.3 Fluorescence detection

After incubation, 100 μL was withdrawn from each tube, mixed with 100 μL of the fluorescence ROSE reagent (an AO/EB-based viability staining reagent, proprietary formulation), and incubated in the dark at room temperature for 5 min. Fluorescence intensity was measured using a SpectraMax iD3 microplate reader (Molecular Devices) at an excitation wavelength of 485 nm and emission wavelengths of 530 nm (green fluorescence, representing viable bacteria) and 620 nm (red fluorescence, representing dead bacteria). Susceptibility was categorized using a two-tier system—susceptible (S) or resistant (R)— based on the fluorescence residual rate (FRR), calculated as follows:

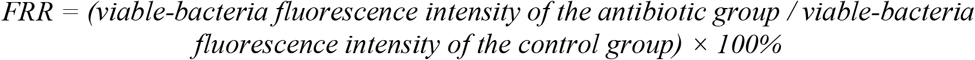

An FRR ≤ 50% was predefined as susceptible (i.e., a ≥50% reduction in viable-bacteria fluorescence after antibiotic exposure relative to control), whereas an FRR > 50% was classified as resistant. This cut-off was established through preliminary optimization experiments using standard strains (see below).

#### 2.2.4 Quality control

Each run included the H. pylori reference strain ATCC 43504 (susceptible control) and a clarithromycin-resistant strain validated at our center. Based on three replicate measurements of five randomly selected specimens, the inter-assay coefficient of variation of the FRR was below 8%.

**Total turnaround time:** approximately 1 hour from biopsy collection to result reporting.

### 2.3 Conventional culture-based susceptibility testing (reference method)

Paired biopsy specimens were processed within 2 hours. After grinding, specimens were inoculated onto H. pylori-selective medium (Columbia agar base supplemented with 7% defibrinated horse blood, vancomycin 10 mg/L, trimethoprim 5 mg/L, polymyxin B 2,500 IU/L, and amphotericin B 2 mg/L) and cultured at 37 °C under microaerobic conditions for 5–7 days. Species identification was performed by Gram staining and catalase, oxidase, and urease tests. Minimum inhibitory concentrations (MICs) were determined using E-test strips (bioMérieux) and interpreted according to the EUCAST v16.1 MIC criteria for H. pylori**Error! Unknown switch argument**.: amoxicillin ≤0.125 μg/mL susceptible and >0.125 μg/mL resistant; clarithromycin ≤0.125 μg/mL susceptible and >0.5 μg/mL resistant; and levofloxacin ≤1.0 μg/mL susceptible and >1.0 μg/mL resistant.

### 2.4 Statistical analysis

The concordance rate was calculated as the proportion of concordant samples (culture-susceptible/ROSE-susceptible plus culture-resistant/ROSE-resistant) among all samples. Sensitivity (the proportion of culture-defined resistant strains correctly identified by the ROSE assay), specificity (the proportion of culture-defined susceptible strains correctly identified by the ROSE assay), positive predictive value (PPV), and negative predictive value (NPV) were calculated, together with their 95% confidence intervals (Wilson score method), using conventional culture (E-test) as the reference standard. Cohen’s kappa coefficient was used to assess agreement beyond chance and was interpreted as follows: ≤0.20, slight; 0.21– 0.40, fair; 0.41–0.60, moderate; 0.61–0.80, substantial; and >0.80, almost perfect agreement^**Error! Unknown switch argument**.^. The 95% confidence intervals of Cohen’s κ coefficients were calculated using the normal approximation method.Statistical analyses were performed using SPSS 26.0 (IBM).

## 3. Results

### 3.1 Baseline patient characteristics

A total of 40 H. pylori-positive patients were enrolled (mean age 43.9 ± 16.2 years; 33 men and 7 women). For all 40 patients, the ROSE and culture-based susceptibility results were compared for three antibiotics—amoxicillin, levofloxacin hydrochloride, and clarithromycin. Fourteen patients were excluded because of unsuccessful culture, leaving 26 patients per antibiotic for statistical analysis.

### 3.2 Concordance between the ROSE and culture methods

Table 1 shows the cross-tabulation of results for each antibiotic. The overall concordance rates of the fluorescence ROSE assay were 84.6% (22/26) for amoxicillin, 76.9% (20/26) for levofloxacin, and 69.2% (18/26) for clarithromycin. Cohen’s kappa coefficients indicated moderate agreement for all three antibiotics (κ = 0.523, 0.539, and 0.412 for amoxicillin, levofloxacin, and clarithromycin, respectively). Sensitivity, specificity, PPV, NPV, and their 95% CIs of the ROSE assay were derived from 2 × 2 tables against the culture gold standard (Table 2). Sensitivity and NPV were both 100.0% for amoxicillin and levofloxacin, and sensitivity was 90.9% for clarithromycin, indicating a low probability that the ROSE assay misses resistant strains and relatively high reliability for clinically ruling out resistance; however, given the limited sample size, these findings require validation in larger cohorts. For specificity, amoxicillin performed best (82.6%), followed by levofloxacin (68.4%) and clarithromycin (53.3%), showing that the ability to distinguish susceptible strains varied among the drugs; the PPVs of all three drugs were moderate, at 42.9%, 53.9%, and 58.8%, respectively.

**Table 1.** Cross-tabulation of the fluorescence ROSE assay versus culture-based susceptibility testing.

| Antibiotic | Culture S /<br>ROSE S | Culture S /<br>ROSE R | Culture R /<br>ROSE S | Culture R /<br>ROSE R | Concordance<br>rate | Kappa (95%<br>CI) |
| --- | --- | --- | --- | --- | --- | --- |
| Amoxicillin | 19 | 4 | 0 | 3 | 84.6% | 0.523 (0.093–<br>0.953) |
| Levofloxacin | 13 | 6 | 0 | 7 | 76.9% | 0.539 (0.215–<br>0.862) |
| Clarithromycin | 8 | 7 | 1 | 10 | 69.2% | 0.412 (0.074–<br>0.751) |

Note: S, susceptible; R, resistant. Concordance rate = (culture S/ROSE S + culture R/ROSE R)/total. Cohen's $\kappa$ was interpreted as $\leq 0.20$ slight, 0.21–0.40 fair, 0.41–0.60 moderate, 0.61–0.80 substantial, and $> 0.80$ almost perfect agreement. The 95% confidence intervals of $\kappa$ were calculated by the normal approximation method.

**Table 2.** Diagnostic metrics of the fluorescence ROSE susceptibility assay.

| Antibiotic | Sensitivity, % (95% CI) | Specificity, % (95% CI) | PPV, % (95% CI) | NPV, % (95% CI) |
| --- | --- | --- | --- | --- |
| Amoxicillin | 100.0 (43.9–100.0) | 82.6 (62.9–93.0) | 42.9 (15.8–74.9) | 100.0 (83.2–100.0) |
| Levofloxacin | 100.0 (64.6–100.0) | 68.4 (46.0–84.6) | 53.9 (29.1–76.8) | 100.0 (77.2–100.0) |
| Clarithromycin | 90.9 (62.3–98.4) | 53.3 (30.1–75.2) | 58.8 (36.0–78.4) | 88.9 (56.5–98.0) |

Note: 95% confidence intervals were calculated by the Wilson score method. PPV, positive predictive value; NPV, negative predictive value.

### 3.3 Analysis of discordant results

For amoxicillin, four results were discordant, all of which were ROSE false positives (ROSE resistant, culture susceptible), with no false negatives. The absence of false negatives indicates that the ROSE assay detected all culture-confirmed resistant strains, with a very low risk of missed detection, corresponding to a sensitivity of 100%; the four false positives reduced specificity to 82.6%. Mechanistically, amoxicillin is a β-lactam antibiotic that acts only on the cell wall of proliferating bacteria; some H. pylori within biopsy specimens grow slowly or are metabolically quiescent, so cell-wall inhibition was not fully manifested after 1 h of drug incubation, and the FRR did not reach the susceptibility threshold, producing false positives. In addition, variation in bacterial load among biopsy specimens alters the effective drug concentration in the well, which can likewise interfere with interpretation.

Among the three antibiotics, clarithromycin had the most discordant cases (eight in total), predominantly false positives: seven were ROSE-resistant but culture-susceptible, and only one was a false negative (ROSE-susceptible but culture-confirmed resistant). Clarithromycin targets the bacterial 50S ribosome, inhibiting protein synthesis without directly damaging the cell membrane, whereas the fluorescence assay used in this study relies on cell-membrane metabolic activity for its readout. After 1 h of drug treatment, susceptible strains with low MICs showed a membrane fluorescence decrease of less than 50%, which was readily misclassified as resistant; this is an inherent methodological limitation of this fluorescence system when evaluating macrolides. The single false negative was likely related to the 23S rRNA A2142G/A2143G resistance mutation^**Error! Unknown switch argument**.^: this mutation imposes a fitness cost, so the strain has low baseline metabolic activity in the absence of drug and spontaneously exhibits a reduced fluorescence signal, leading to misclassification. These results suggest that the current 50% FRR threshold readily produces clarithromycin false positives and that modestly raising the susceptibility threshold (e.g., defining FRR ≤ 60% as susceptible) could improve specificity.

For levofloxacin, six results were discordant, all false positives with no false negatives; consistent with amoxicillin, this indicates that the ROSE assay detected all culture-resistant strains, with little risk of missed detection. Levofloxacin targets DNA gyrase and exerts its antibacterial effect through DNA breakage, but the metabolic inhibition triggered by DNA damage is difficult to fully manifest within a short 1 h incubation; some low-MIC susceptible strains retained relatively high cell viability after treatment at the breakpoint concentration, with FRR > 50%, ultimately yielding false positives.

### 3.4 Turnaround time and cost

The fluorescence ROSE assay reported results within 1 hour (range 55–65 min), whereas conventional culture required 5–7 days (median 6 days). The preliminary estimated cost saving was 3,000–5,000 CNY per patient, mainly attributable to the elimination of culture media, prolonged occupation of incubation equipment, and repeat clinic visits to collect reports.

Using amoxicillin as an example, representative fluorescence ROSE images and quantitative readouts before and after 1 h of antibiotic exposure are shown below.

A. **Susceptible strain (amoxicillin):** after exposure (Fig. 3B), fluorescence intensity decreased markedly relative to before exposure (Fig. 3A) (FRR ≤ 50%), indicating metabolic inhibition. Green fluorescence (viable bacteria) was markedly attenuated, with a corresponding increase in red fluorescence (non-viable/dead bacteria).
B. **Resistant strain (amoxicillin):** fluorescence intensity showed no significant change (FRR > 50%), indicating that bacterial metabolic activity was maintained even under antibiotic exposure.
C. Fluorescence residual rates of the three antibiotics, stratified by culture-based susceptibility status. The horizontal dashed line indicates the 50% susceptibility threshold. Error bars represent the standard deviation of three replicate measurements.

**Figure 1.**
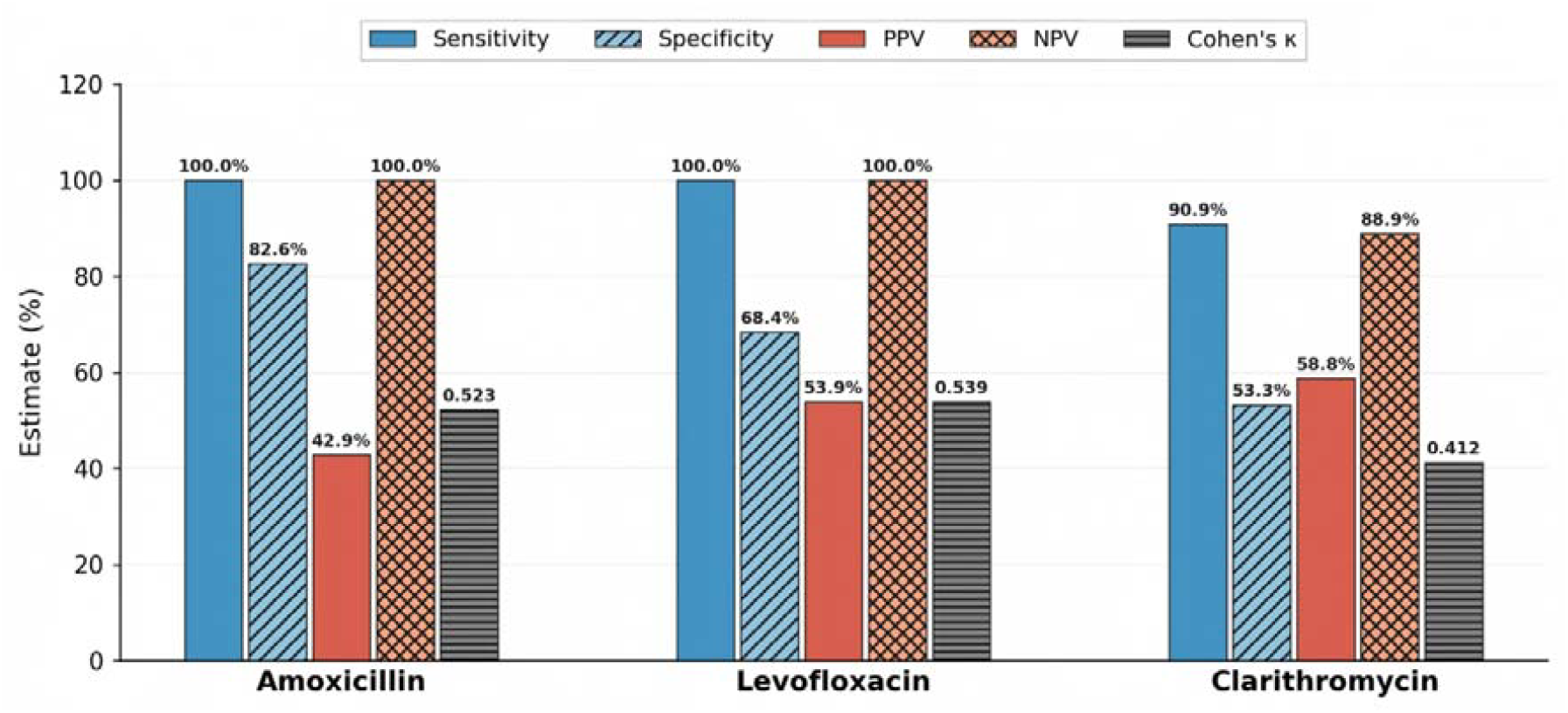
Comparison of diagnostic performance metrics (grouped bar chart) Note: Cohen’s kappa coefficient was multiplied by 100 for display on the same axis as the other metrics (the values labeled in the figure are the original κ values, without a percent sign).

**Figure 2.**
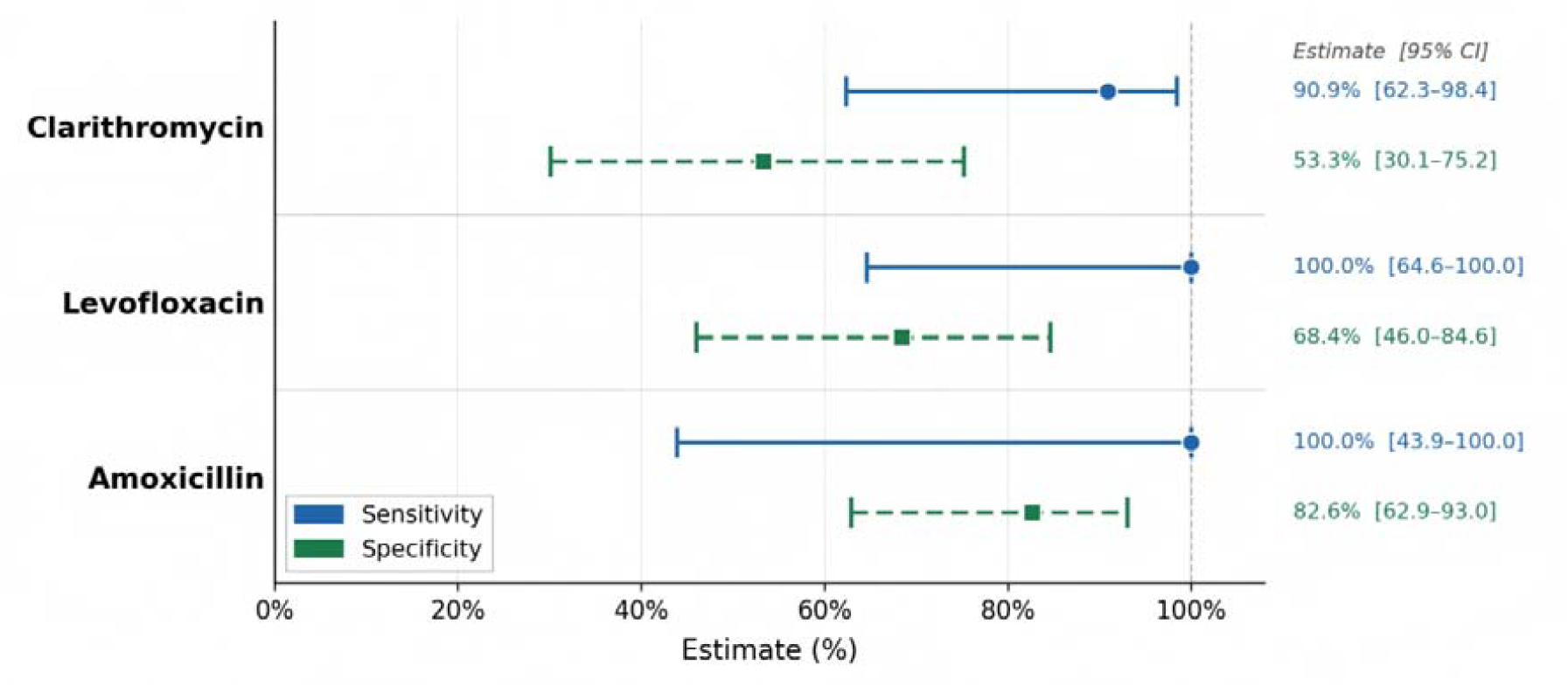
Forest plot comparing the sensitivity and specificity of the three antibiotics (with 95% CIs) Note: 95% confidence intervals were calculated by the Wilson score method.

**Figure 3.**
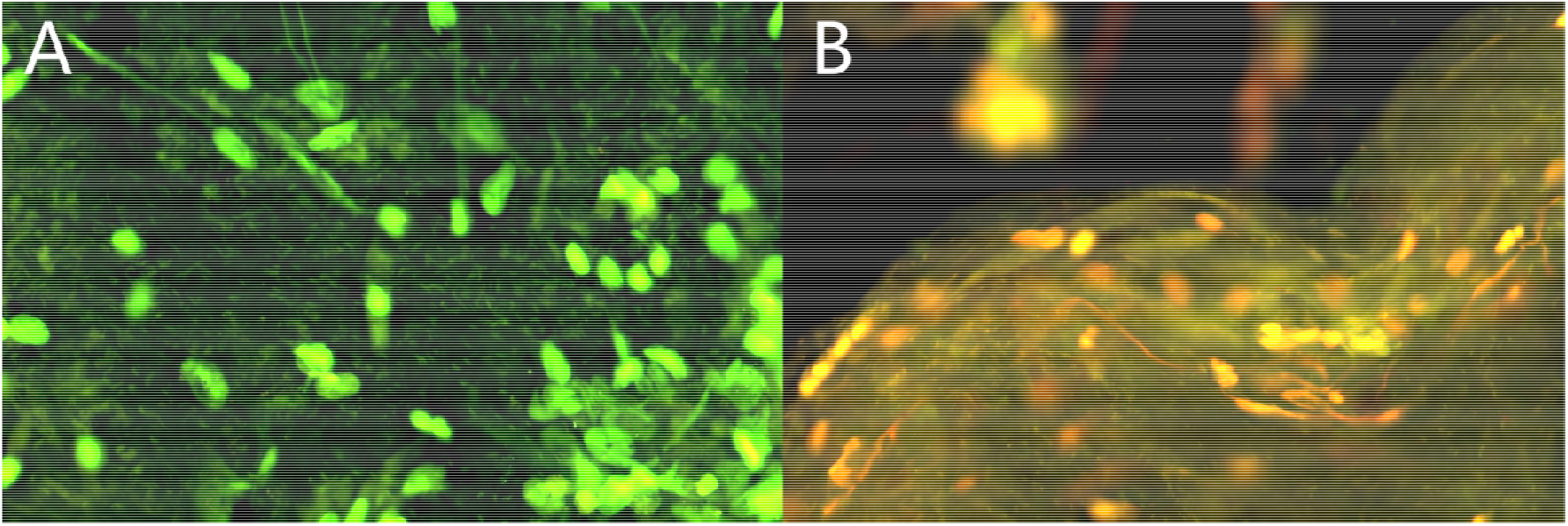
Note: A, amoxicillin-susceptible strain before antibiotic exposure (400×); B, amoxicillin-susceptible strain after antibiotic exposure (400×). Green fluorescence (excitation 485 nm/emission 530 nm) represents viable bacteria, and red fluorescence (emission 620 nm) represents dead bacteria.

**Figure 4.**
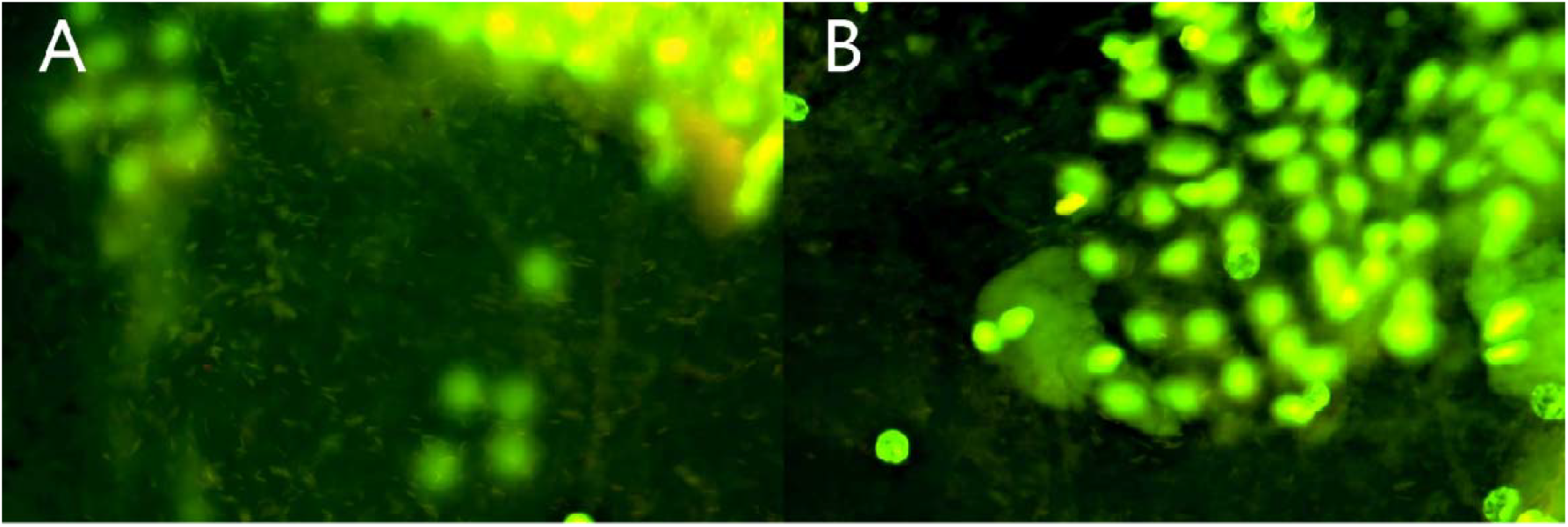
Note: A, amoxicillin-resistant strain before antibiotic exposure (400×); B, amoxicillin-resistant strain after antibiotic exposure (400×).

**Figure 5.**
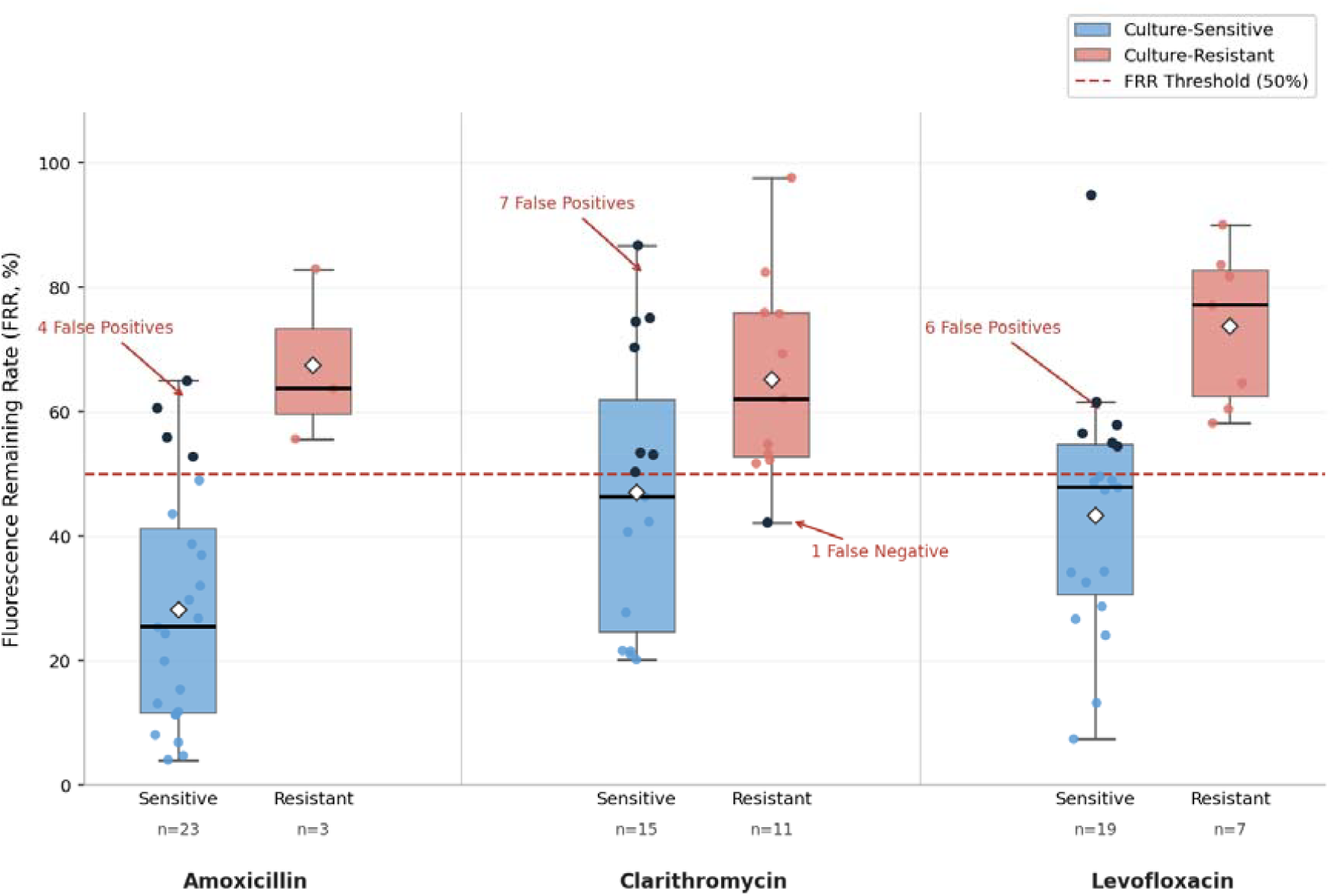
Note: Fluorescence residual rate (FRR) stratified by culture-based susceptibility status; error bars represent the standard deviation of three replicates. Scale bar = 20 μm. FRR, fluorescence residual rate. Black scatter points indicate discordant results (false positives/false negatives).

## 4. Discussion

### 4.1 Main findings and per-antibiotic performance

To our knowledge, this is the first report of a culture-free rapid phenotypic susceptibility test for H. pylori that can use gastric mucosal biopsy specimens directly and yield results within 1 hour. Since the standardization of phenotypic susceptibility testing in the 1990s, H. pylori AST has relied on bacterial culture^**Error! Unknown switch argument**.^; the 5–7-day testing cycle has perpetuated empirical treatment and may have indirectly promoted the spread of resistance. By detecting antibiotic-induced metabolic inhibition in real time, the fluorescence ROSE assay bypasses bacterial isolation and growth, thereby breaking this pattern.

Diagnostic performance differed among the antibiotics. Amoxicillin had the highest concordance rate (84.6%), with moderate inter-method agreement (κ = 0.523), a sensitivity of 100%, and a relatively high specificity (82.6%), indicating a strong ability to identify resistant strains. Given that there is currently no reliable genotypic alternative for amoxicillin testing, this performance is of considerable clinical importance. Levofloxacin showed moderate agreement (κ = 0.539); although its concordance rate (76.9%) was lower than that of amoxicillin, it likewise showed high sensitivity (100%) and demonstrated preliminary feasibility at the proof-of-concept stage. However, its false-positive rate was relatively high (6/26), suggesting that the current threshold needs to be optimized specifically for levofloxacin. Clarithromycin had the lowest concordance rate (69.2%, κ = 0.412), with discordance driven mainly by false positives and likewise requiring threshold optimization. Considering the discordance data across all three drugs, the discordant results were predominantly false positives, and the current 50% FRR threshold is conservative overall, with a higher risk of overestimating resistance than of missing it. Future work could adjust the fluorescence thresholds for each drug and evaluate improvements in specificity and concordance, while expanding the sample size to verify whether this error pattern is stable.

The pattern observed for clarithromycin warrants specific discussion. The 50% FRR threshold was established using reference strains with clearly distinct susceptible and resistant MICs. However, clarithromycin-resistant strains (mainly carrying 23S rRNA mutations) may retain sufficient overall metabolic activity during the 1 h exposure, preventing the FRR from falling to ≤50% and producing seven false positives. This suggests that a uniform 50% threshold may be too conservative for clarithromycin and prone to generating numerous false positives; appropriately raising the FRR susceptibility threshold (e.g., to 60%) would broaden the susceptible category and reduce false positives. This threshold heterogeneity, rooted in the resistance mechanism rather than in a technical limitation of the ROSE assay itself, indicates that interpretive criteria should be optimized separately for each antibiotic. We are currently developing a machine-learning-based multi-threshold optimization approach that fits fluorescence readouts to MIC distributions; preliminary analyses indicate that this approach can substantially improve interpretive concordance for clarithromycin.

### 4.2 Clinical value of amoxicillin testing

Amoxicillin is the core antibiotic in bismuth quadruple and vonoprazan dual therapies for H. pylori eradication, and strain resistance to it directly causes eradication failure. However, mainstream susceptibility testing methods all have inherent structural deficiencies in detecting amoxicillin resistance, leaving the clinic without a rapid, reliable option. The ROSE fluorescence technique, based on the phenotypic principle of directly detecting bacterial metabolic activity, circumvents the technical shortcomings of these methods and is currently the only novel approach capable of completing accurate amoxicillin susceptibility classification within 1 hour.

#### 4.2.1 Inherent limitations of conventional culture-based susceptibility testing

Agar dilution, E-test, disk diffusion, and broth microdilution are all CLSI-endorsed phenotypic gold standards and can, in principle, quantitatively determine the MICs of various antibiotics. However, when applied to amoxicillin resistance in H. pylori, their inter-method agreement and accuracy are markedly lower than for clarithromycin, levofloxacin, metronidazole, and other agents.

i. **Poor inter-method agreement, with prominent missed detection and false resistance**. In a 2020 ROC analysis of 72 Indonesian clinical H. pylori isolates by Miftahussurur et al., E-test detection of amoxicillin resistance had a sensitivity of only 33.3%, a specificity of 95.7%, and an AUC of 0.783, with a kappa of only 0.25 versus agar dilution, indicating substantial systematic bias; in the same batch, E-test sensitivities for levofloxacin and metronidazole reached 88.9% and 86.7%, respectively^**Error! Unknown switch argument**.^. A study of 77 strains by Ogata et al. further showed that the correlation coefficient between disk diffusion and agar dilution for the amoxicillin MIC was r = −0.36 (p = 0.0015), the worst among all antibiotics tested; the amoxicillin resistance rate determined by disk diffusion was as high as 68.8%, versus only 10.4% by the agar-dilution gold standard—an extremely high false-resistance rate**Error! Unknown switch argument**..
ii. **Underlying mechanisms of measurement bias and practical shortcomings**. H. pylori can develop amoxicillin resistance through multiple routes, including β-lactamase hydrolysis, reduced penicillin-binding protein (PBP) affinity, and activation of active efflux pumps^**Error! Unknown switch argument**.^. Five independent and additive resistance pathways have been identified (Table 3), with no single marker gene covering the entire resistance phenotype:

**Table 3.**
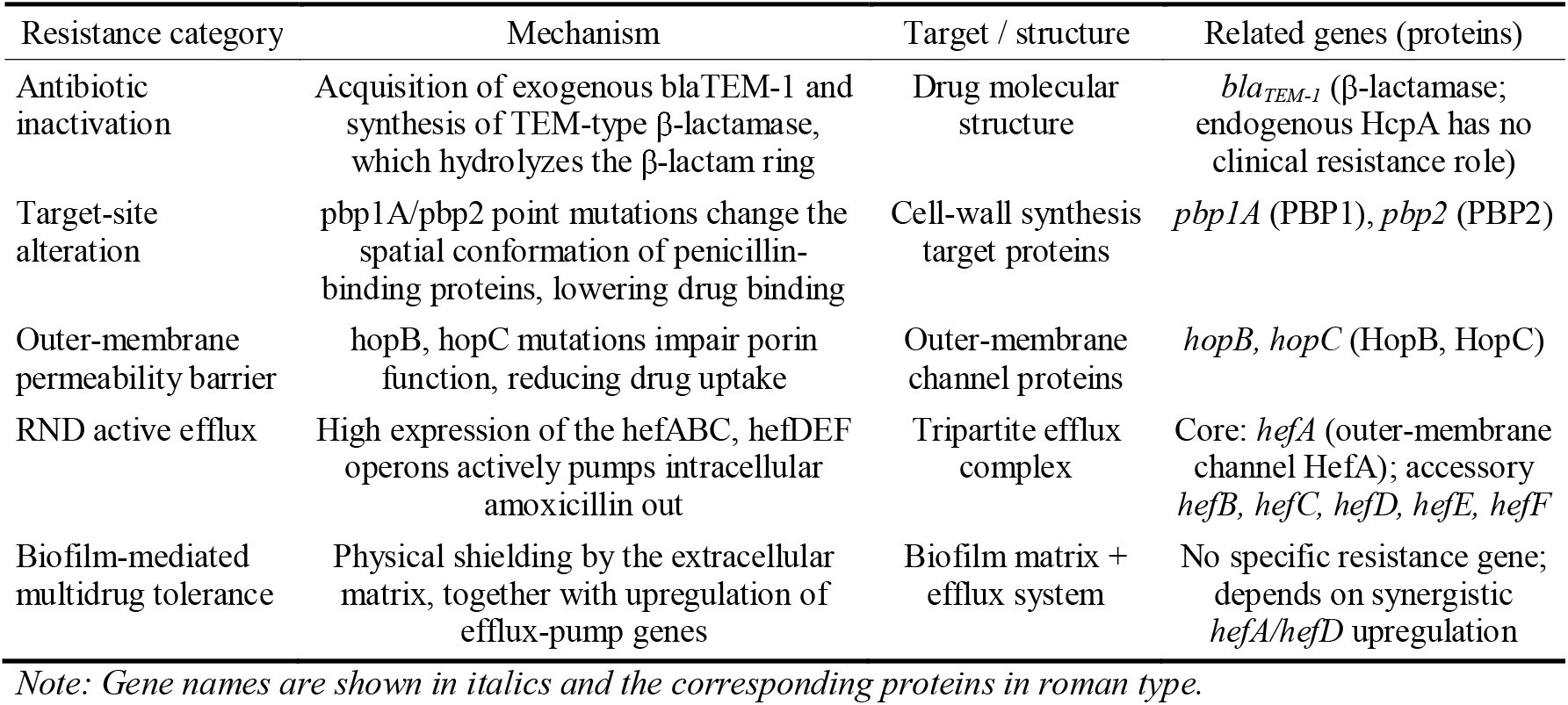
Summary of amoxicillin resistance mechanisms in H. pylori.

| Resistance category | Mechanism | Target / structure | Related genes (proteins) |
| --- | --- | --- | --- |
| Antibiotic inactivation | Acquisition of exogenous *bla*<sub>TEM-1</sub> and synthesis of TEM-type β-lactamase, which hydrolyzes the β-lactam ring | Drug molecular structure | *bla*<sub>TEM-1</sub> (β-lactamase; endogenous HcpA has no clinical resistance role) |
| Target-site alteration | *pbp1A/pbp2* point mutations change the spatial conformation of penicillin-binding proteins, lowering drug binding | Cell-wall synthesis target proteins | *pbp1A* (PBP1), *pbp2* (PBP2) |
| Outer-membrane permeability barrier | *hopB*, *hopC* mutations impair porin function, reducing drug uptake | Outer-membrane channel proteins | *hopB*, *hopC* (HopB, HopC) |
| RND active efflux | High expression of the *hefABC*, *hefDEF* operons actively pumps intracellular amoxicillin out | Tripartite efflux complex | Core: *hefA* (outer-membrane channel HefA); accessory *hefB*, *hefC*, *hefD*, *hefE*, *hefF* |
| Biofilm-mediated multidrug tolerance | Physical shielding by the extracellular matrix, together with upregulation of efflux-pump genes | Biofilm matrix + efflux system | No specific resistance gene; depends on synergistic *hefA/hefD* upregulation |

Note: Gene names are shown in italics and the corresponding proteins in roman type.

1. Antibiotic inactivation: strains acquire exogenous *bla*_*TEM-1*_ and express TEM-type β-lactamase, which hydrolyzes the β-lactam ring of amoxicillin, mediating high-level resistance**Error! Unknown switch argument**.;
2. Target protein structural alteration: *pbp1A* and *pbp2* point mutations alter the drug-binding regions of PBP1/PBP2, producing stable, heritable low-to-moderate resistance^[12,13]^;
3. Outer-membrane permeability barrier: *hopB* and *hopC* mutations impair outer-membrane porin function, markedly reducing drug uptake^[14]^;
4. RND-type active efflux: overexpression of the *hefABC* and *hefDEF* operons actively expels intracellular amoxicillin through the HefA outer-membrane channel protein^[15-17]^;
5. Biofilm-mediated reversible tolerance: physical shielding by the extracellular matrix, combined with upregulation of the *hefA*/*hefD* efflux genes, doubly weakens bactericidal efficacy, with no specific gene marker^[18]^.

The five resistance pathways described above either depend on independent genetic markers (e.g., *bla*_*TEM-1*_, *pbp1A*) or leave no genetic-mutation imprint at all (e.g., biofilm-mediated efflux-pump overexpression), so no single-gene test can cover the full spectrum. In contrast, the ROSE fluorescence method directly quantifies the metabolic activity of viable bacteria after antibiotic exposure; regardless of which pathway confers resistance, as long as the drug fails to effectively inhibit metabolism, resistance is captured—this is precisely why the method inherently bypasses all of the above pathways in principle.

Moreover, H. pylori requires a microaerobic environment of 5% O□ and 10% CO□ and a culture cycle of up to 5–7 days, imposing high demands on the equipment and technical capacity of primary-care institutions; before sampling, proton pump inhibitors must be withheld for 2 weeks and antibiotics for 4 weeks, and poor patient adherence leaves many endoscopic samples without valid susceptibility results^[10]^.

#### 4.2.2 Structural blind spots of molecular testing

PCR-derived techniques, first-generation sequencing and whole-genome sequencing (WGS), FISH, and gene chips are mature tools for the rapid screening of clarithromycin and levofloxacin resistance, on the premise that the resistance mechanisms of these two drug classes are simple, with fixed mutation sites—clarithromycin resistance arises almost entirely from the 23S rRNA A2142G/A2143G mutations, and levofloxacin resistance is concentrated in high-frequency mutations such as N87K and D91N in the *gyrA* QRDR. The commercial GenoType HelicoDR kit detects clarithromycin resistance with a sensitivity and specificity of 94% and 99%, respectively^[19]^. However, this technical route is not suitable for the accurate prediction of amoxicillin resistance.

i. **Large-sample clinical evidence: genotype cannot predict the true resistance phenotype**. A retrospective study in Nanjing reported by Wang et al. in 2026 included a genotype–phenotype matching analysis of 7,227 patients (5,729 culture-positive) from 2018– 2024; the kappa coefficients for clarithromycin 23S rRNA and levofloxacin *gyrA* reached 0.9046 and 0.9187, respectively, indicating excellent agreement, whereas that for amoxicillin was only 0.0239, and those for tetracycline, furazolidone, and metronidazole were likewise close to zero, indicating that testing *pbp1A* mutations alone cannot reflect the true resistance status^[20]^. A WGS analysis of 141 clinical strains from Hangzhou by Wang P et al. likewise showed that the concordance of amoxicillin resistance prediction based on *pbp1A* mutations was only 83.69%, lower than that for clarithromycin (92.91%) and levofloxacin (89.13%)^[21]^. WGS can detect only mutations already in the database and cannot identify emerging mutations or non-genetically mediated resistance mechanisms such as biofilm and efflux pumps, readily producing false-susceptible results^[10,21]^.
ii. **Common limitations of existing molecular platforms**. The various PCR-derived assays (PCR-RFLP, DPO-PCR, qPCR, ddPCR) can detect only known resistance mutations already reported in databases; fluorescence in situ hybridization (FISH) lacks a mature commercial probe system for amoxicillin resistance because the target genes are dispersed and lack a unified target sequence (unlike clarithromycin 23S rRNA, for which such probes exist^[22]^); and resistance gene chips are limited to the sites entered at design and are costly to make comprehensive. More critically, the reversible high-level expression of biofilm barriers and efflux pumps produces drug tolerance only at the phenotypic level, without any mutation marker in the gene sequence, so all gene-based detection methods have an inherent blind spot^[18,20]^. On the basis of these deficiencies, the Maastricht VI/Florence consensus notes that guiding amoxicillin-containing eradication regimens by genotypic testing alone has clear clinical limitations^[3,10]^.

#### 4.2.3 Clinical advantages of the ROSE fluorescence technique

i. **Core principle: mechanism-independent direct phenotypic detection**. The ROSE fluorescence technique does not rely on resistance-gene identification but directly quantifies the viable-bacteria fluorescence residual rate of H. pylori after amoxicillin exposure. Regardless of whether a strain develops resistance through any single or combined mechanism—β-lactamase, PBP mutation, porin deficiency, efflux pump, or biofilm—as long as the metabolic activity of viable bacteria is not inhibited after drug intervention (FRR ratio ≥ the 50% threshold), it can be accurately identified, thereby circumventing the genetic blind spots of molecular methods^[10]^. Its interpretive logic is consistent with the agar-dilution gold standard, but it compresses the 5–7-day cycle to 1 hour, making it compatible with the needs of point-of-care endoscopic diagnosis and treatment.
ii. **Comparison of diagnostic performance**. Conventional methods have marked performance deficits: E-test sensitivity is only 33.3%, missing nearly two-thirds of resistant strains^[8]^, and genotypic testing has a kappa of only 0.0239, with predictions approaching random^[20]^. By contrast, in 26 clinical samples the ROSE fluorescence technique achieved a resistance-detection sensitivity of 100.0% (43.9%–100.0%) and a specificity of 82.6% (62.9%–93.0%), with a 100% test-success rate. Thus, the ROSE fluorescence technique is more sensitive than the E-test and, in principle, resolves the structural genotype–phenotype disconnect of molecular testing. This three-way comparison clearly shows that, in amoxicillin susceptibility testing, the E-test carries a nearly two-thirds risk of missing resistance, genotypic methods achieve near-random agreement, and the present ROSE method achieved zero missed detection (sensitivity 100.0%), confirming the clinical feasibility of the direct phenotypic testing route.
iii. **Clinical application value**. On the basis of the above principle advantage of direct phenotypic detection and the preliminary validation data, the ROSE fluorescence method provides a feasible laboratory basis for the accurate use of amoxicillin in bismuth quadruple and vonoprazan regimens and holds promise for completing same-day susceptibility testing concurrently with endoscopic diagnosis and treatment, thereby moving H. pylori eradication from empirical toward individualized therapy.

### 4.3 Limitations and future directions

This study has several limitations. It is a small-sample, single-center preliminary study intended to establish proof of concept. The predefined 50% threshold has not been optimized and very likely needs to be set differently for different antibiotics; because the present analysis used a uniform threshold, it represents a “worst-case” scenario. The reference method (culture-based susceptibility testing) itself has inherent limitations: some resistant strains may carry a fitness cost^[7]^, leading to culture failure or delayed growth and thus missed detection, and culture conditions (microaerobic atmosphere, medium composition) may differ from the in vivo environment. In addition, this study did not evaluate the effect of mixed infection or heteroresistance on the fluorescence readout.

To address these limitations and further translate this technology into clinical practice, we have initiated a prospective randomized controlled trial comparing eradication therapy guided by the 1 h fluorescence ROSE assay with empirical bismuth quadruple therapy for H. pylori (registration no. [ChiCTR2600125137]). The primary endpoint is the eradication rate; secondary endpoints include antibiotic consumption, adverse events, and cost-effectiveness. This trial will provide the highest level of evidence for the application of rapid phenotypic susceptibility testing in the clinical diagnosis and treatment of H. pylori.

In the broader field of H. pylori diagnostics, our research team has established a systematic research pathway: from high-accuracy diagnosis (approximately 10 prior studies based on fluorescence ROSE), to rapid phenotypic susceptibility testing (the present study), to clinical endpoint validation (the ongoing randomized controlled trial). This integrated pathway, built around a single fluorescence detection platform, forms a closed loop from diagnosis to susceptibility testing to clinical validation, offering a new approach to methodological innovation in gastroenterology.

## 5. Conclusions

This study preliminarily confirms the feasibility of culture-free 1-hour phenotypic susceptibility testing for H. pylori using fluorescence rapid on-site evaluation technology. The method compresses conventional phenotypic susceptibility testing—which requires 5–7 days—to within 1 hour, providing a laboratory basis for formulating susceptibility-guided individualized regimens on the same day. It is particularly valuable for amoxicillin testing, for which no genotypic alternative currently exists. With ongoing algorithm optimization and the randomized controlled trial underway, the culture-free 1-hour nature of this technology demonstrates a clear path toward clinical translation.

## Acknowledgments

The authors dedicate this work to the entire research team and thank them for their sustained efforts throughout the process of advancing this technology from concept to clinical application.

## Conflict of interest statement

The authors declare no relevant conflicts of interest.

## Notes

### Competing Interest Statement

The authors have declared no competing interest.

